# Altered hepatic metabolism mediates sepsis preventive effects of reduced glucose supply in infected preterm newborns

**DOI:** 10.1101/2023.09.11.557116

**Authors:** Ole Bæk, Tik Muk, Ziyuan Wu, Yongxin Ye, Bekzod Khakimov, Alessandra Maria Casano, Bagirath Gangadharan, Ivan Bilic, Anders Brunse, Per Torp Sangild, Duc Ninh Nguyen

**Author notes:** Corresponding author: Duc Ninh Nguyen, Section for Comparative Pediatrics and Nutrition, Department of Veterinary and Animal Sciences, University of Copenhagen, Dyrlægevej 68, DK-1870 Frederiksberg C, Denmark. Tel: +45. 35333250. Co-first authors (Ole Bæk, Tik Muk). At time of study. Current affiliation: independent consultant.

## Abstract

Preterm infants are susceptible to neonatal sepsis, a syndrome of pro-inflammatory activity, organ damage and altered metabolism following infection. Given the unique metabolic challenges and poor glucose regulatory capacity of preterm infants, their glucose intake during infection may have a high impact on the degree metabolism dysregulation and organ damage. Using a preterm pig model of neonatal sepsis, we previously showed that a drastic restriction in glucose supply during infection protects against sepsis via suppression of glycolysis-induced inflammation, but results in severe hypoglycemia. Now we explored clinically relevant options of reducing glucose intake to decrease sepsis risk, without causing hypoglycemia and further explore the involvement of the liver in these protective effects. We found that a reduced glucose regime during infection increased survival via reduced pro-inflammatory response, while maintaining normoglycemia. Mechanistically, this intervention enhanced hepatic oxidative phosphorylation and possibly gluconeogenesis, and dampened both circulating and hepatic inflammation. However, switching from a high to a reduced glucose supply after debut of clinical symptoms did not prevent sepsis, suggesting metabolic conditions at the start of infection are key in driving the outcome. Finally, an early therapy with purified human inter-alpha inhibitor protein, a liver derived anti-inflammatory protein, partially reversed the effects of low parenteral glucose provision, likely by inhibiting neutrophil functions that mediate pathogen clearance.

Our findings suggest a clinically relevant regime of reduced glucose supply for infected preterm infants could prevent or delay the development of sepsis in vulnerable neonates.

## Introduction

Preterm infants (born before 37 weeks of gestation) are particularly susceptible to serious neonatal infections potentially leading to sepsis, a state of life-threatening organ dysfunction^1^. Most episodes of late-onset sepsis in these infants (occurring >48 hours after birth) are caused by coagulase negative staphylococci (CONS), such as *Staphylococcus epidermidis*^2,3^, which may enter the circulation via indwelling medical devices or from the gastrointestinal tract^4,5^. The development of sepsis is often attributed to the immature immune systems in preterm infants, but a recent concept of immunometabolism suggests it may partly be caused by the distinct newborn metabolic state, affected by their limited energy reservoirs^6^. Upon exogenous challenge, immune cells and many other cell types in healthy adults undergo metabolic shift from oxidative phosphorylation (OXPHOS) to glycolysis, producing large amounts of adenosine triphosphate (ATP) to fuel inflammatory responses, facilitating efficient reduction of pathogen burden through immunological resistance^6,7^. However, immune cells can also employ a tolerance strategy, limiting the active resistance response and associated immunopathology, via an opposite shift from glycolysis to OXPHOS^6^. This tolerance strategy is found in newborns, especially those born prematurely, as they have low energy stores, but high demands for growth and development. Therefore, energy tends to be prioritized for vital organ functions rather than immune responses to external stimuli, explaining why they display low immune cell metabolism^6,8^ and can withstand 10-100 times greater pathogen loads than adults before showing clinical symptoms^9^.

The liver plays a critical role during bloodstream infections, being a well perfused organ with resident immune cells. Besides its role in producing important immunomodulatory plasma proteins, the role of hepatic metabolism during infections has recently emerged^9^. For instance, blocking hepatic glycolysis lowers overall energy expenditure and improves OXPHOS, which facilitates immune tolerance and better survival during polymicrobial sepsis in mice^10^. This suggests that manipulation of hepatic energy metabolism may affect the systemic immune responses and be a promising avenue to reduce morbidity during neonatal infections. Progression of sepsis is also concomitant with activation of neutrophils, may lead to increased activity of neutrophil-derived serine proteases that play a key role in sepsis-related tissue damage^11^. Their inhibition has likewise been proposed as a therapeutic strategy to resolve septic conditions^12^. Recent studies have shown that the components of one of the most abundant plasma protein complexes, Inter-Alpha Inhibitor Protein (IAIP), are differentially regulated during sepsis leading to a significant reduction of the whole complex both in adults^13–15^ and neonates^16,17^. Administration of enriched human plasma-derived IAIP (hIAIP) in a murine model of neonatal sepsis has been shown to reduce disease severity and mortality through suppression of pro-inflammatory responses^18^.

Due to the lack of mother’s milk and immature gut, preterm infants often receive glucose-rich parenteral nutrition (PN) during the first few weeks of life to ensure sufficient nutrition until full enteral nutrition can be established. The glucose levels are tailored to provide energy for growth and to avoid hypoglycemia-induced brain injury^19–22^. However, this practice leads to hyperglycemia in up to 80% of very preterm infants during early life^23^. Prolonged glucose-rich PN is related to longer hospitalization in septic infants^21^, which suggests it may affect the blood immune-metabolic axis or hepatic metabolism during infection, facilitating glycolysis and active immune resistance. This is in line with a recent study showing that limited nutrition in the acute phase of bacterial infection in rodents reduced glycolysis and enhanced disease tolerance, leading to better survival^24^. Of note, there are no specific guidelines for parenteral glucose intake during serious neonatal infections.

Preterm pigs provide a unique, prematurely born, animal model, enabling PN via an umbilical catheter and mimicking clinical and cellular infection responses to *S. epidermidis*^25,26^, as seen in septic preterm infants similarly infected with CONS^2^. With this model, we have previously shown that the high parenteral glucose supply used in neonatal intensive care units exaggerated systemic glycolysis, triggering hyper-inflammatory responses and clinical sepsis signs, whereas strict glucose restriction prevented sepsis, but caused hypoglycemia^27^. Here, we further exploited this model to explore how glucose regimes affect hepatic metabolism and glucose homeostasis to both prevent hypoglycemia and to alleviate the risk of sepsis in CONS infected preterm newborns. We identified a reduced parenteral glucose regimen that during neonatal infection both maintained normoglycemia and modulated the hepatic immune-metabolic networks in a manner that avoided excessive energy used for immune responses, leading to improved overall survival. However, postponing the reduction in glucose supply until clinical symptoms started to manifest did not affect clinical deterioration, suggesting metabolic constraints early in the immune response are critical for determining the fate. Lastly, to further explore potential optimizations of the treatment regimen, we evaluated concomitant treatment of pre-term pigs with reduced PN glucose regimen supplemented with intravenous administration of purified hIAIP. In contrast to our original hypothesis, hIAIP supplementation reversed the effects of glucose reduction in this model of neonatal sepsis and provided no survival benefit.

In all, these clinically relevant findings suggest that reduced parenteral glucose supply, together with antibiotics and other supporting care, as a concomitant therapy for CONS infected preterm infants may prevent or delay development of neonatal sepsis.

## Results

### Reduced glucose supply improves glucose homeostasis and survival during neonatal infection

Using preterm pigs as models for infected preterm infants, we have previously shown that hyperglycemia induced by a high parenteral glucose supply (High, 21%, 30g/kg/day), predisposed to sepsis and high mortality following *S. epidermidis* infection. However, a very restrictive glucose supply, below the recommended guidelines for preterm infants^28^ (1.4%, 2g/kg/day) protected against development of sepsis, but caused severe hypoglycaemia^27^.

Here we investigated whether it was feasible to reduce parenteral glucose supply during neonatal infection to an extent that reduced the severity of symptoms, while avoiding hypoglycemia. Preterm pigs were delivered by caesarian section at 90% gestation and nourished with PN using either the high or a reduced glucose (Low, 5%, 7.2 g/kg/day) regimen^28^. The animals were systemically infused with either live *S. epidermidis* (SE, 10^9^ colony forming units (CFU)/kg) or control saline (CON), see Figure 1A for study overview). Animals were then closely monitored for 22 hours, and those showing severe septic symptoms (deep lethargy, hypoperfusion, and blood pH < 7.1 as signs of severe acidosis and impending respiratory/circulatory collapse) were euthanized according to predefined humane endpoints.

**Figure 1:**
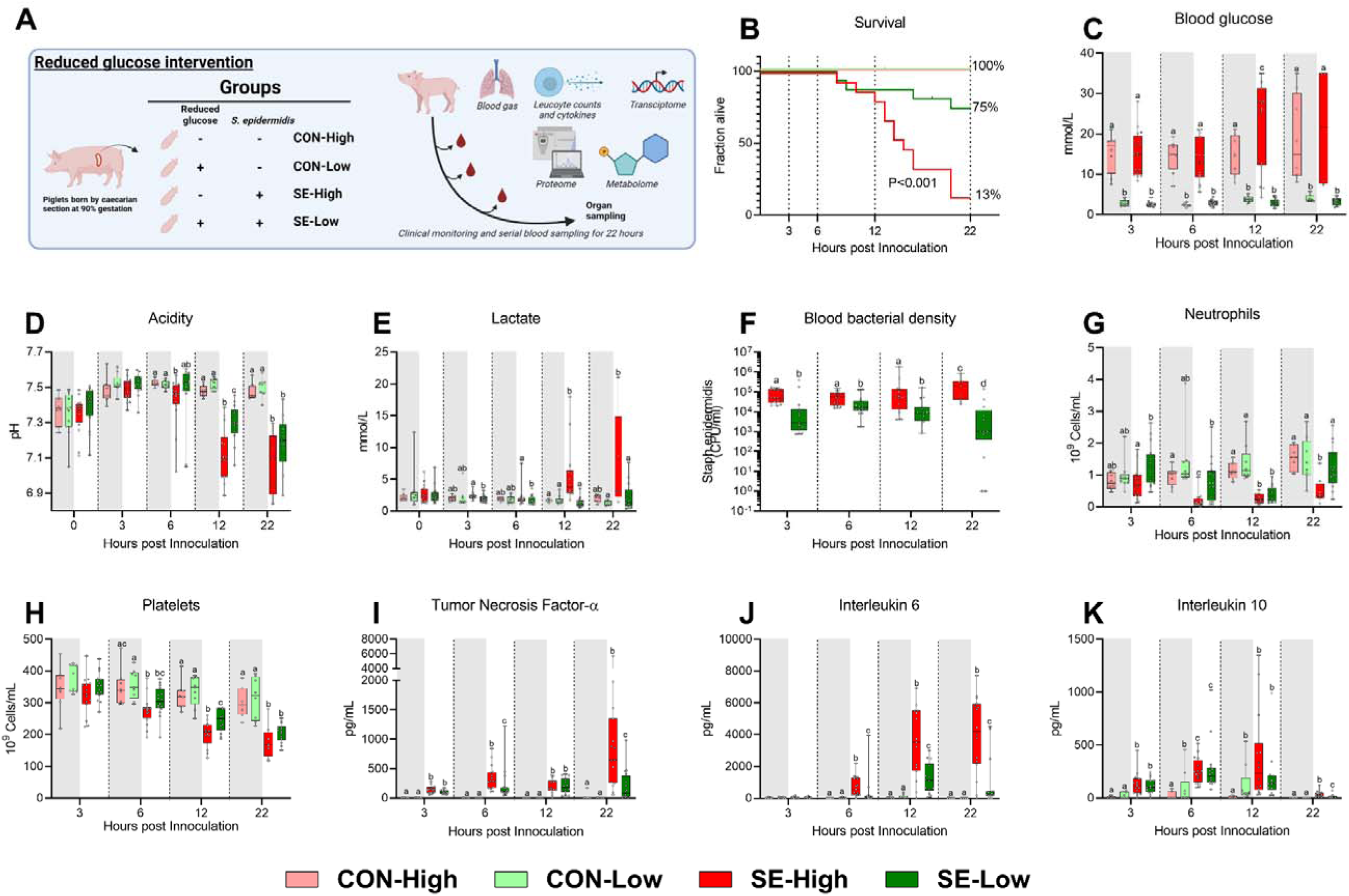
Animal study overview, clinical and immunological results. **A:** Study overview, preterm pigs were nourished with parenteral nutrition (PN) with either high (21%, 30 g/kg/d) or low (5%, 7.2 g/kg/d) glucose supply and infused with either *Staphylococcus epidermidis* or control saline. Animals were followed for 22 hours and blood samples collected for further analysis. Figure created with BioRender.com. **B:** Survival of animals during experiment, presented as time to euthanasia according to predefined humane endpoints with corresponding log-rank test comparing SE-High and SE-Low **C:** Blood glucose, measured by glucose meter 3, 6 and 12 hours after SE inoculation as well as at euthanasia, presented as 95% box plots. **D,E:** Blood gas parameters collected before inoculation with SE (0 hours) and 3, 6 and 12 hours after as well as at euthanasia, presented as 95% box plots. **F:** Blood bacterial density in infected groups at 3, 6, 12 and 22 hours after bacterial inoculation, presented as 95% box plots on a logarithmic scale. **G-K:** Blood hematology and plasma cytokines at 3, 6, 12 and 22 hours after bacterial inoculation, presented as 95% box plots. **C-K:** Data at each time point analyzed separately, bars labeled with different letters are significantly different from each other (P < 0.05), n = 8-9 for control animals, and 10-16 for infected groups.

Reduced glucose supply markedly improved survival following infection, with an almost six-fold reduction in the absolute risk of death (P < 0.001, Figure 1B). The reduced glucose supply expectedly led to lower blood glucose levels, compared to SE-High animals, and blood glucose levels remained stable around 2-3 mmol/l during the whole experiment (Figure 1C). Blood pH started to decrease at a slower rate from 6h onwards in SE-Low animals with correspondingly lower lactate levels, indicating slower rates of glycolytic breakdown of pyruvate (Figure 1D,E). Of note, base excess was improved in SE-Low animals already from 3 h after start of the infection, whereas glucose supply during infection had no impact on partial carbon dioxide pressure (Sup Figure S1 A,B). This indicates that the severity of the metabolic acidosis occurring during neonatal sepsis was alleviated by the reduced glucose regimen. Importantly, despite the marked difference in blood glucose levels, blood osmolality did not differ much among the groups (Sup Figure S1C), underlining that the differences in clinical responses were not driven by a hyperosmolar state induced by the high glucose supply. As such, it appeared that reduction in parenteral glucose supply improved survival and clinical symptoms while still maintaining normoglycemia, in the lower range.

### Glucose homeostasis affects cellular responses to neonatal infection

With a clear clinical benefit of reduced glucose supply observed, we then further investigated the potential underlying mechanisms by measuring the immune cell and molecular responses to the infection. SE-Low animals showed consistently lower blood bacterial densities for the duration of the experiment (Figure 1F). All the main blood leucocyte subsets dropped in response to the infection (Sup Figure S1D, E), but depletions in circulating neutrophils were more pronounced in SE-High vs. SE-Low animals, evident at 6 h post-infection which were only replenished in SE-Low animals (Figure 1G), suggesting either better bone marrow capacity or less neutrophil activation, although this was not further investigated. Also, SE-Low animals showed less severe thrombocytopenia (Figure 1H), and depletions of these blood cellular subsets are known to be associated with increased severity in neonatal sepsis^29–31^.

In parallel, it was evident that the inflammatory response in infected animals with a reduced glucose supply also was dramatically attenuated, with lower levels of plasma TNF-α, IL-6, and IL-10, compared to animals with high glucose supply (Figure 1I-K). Differences in TNF-α and IL-6 levels were already evident at 6 h after inoculation. Interestingly, among uninfected control animals, a reduced glucose supply also increased IL-10 levels (Figure 1K).

This indicates that a reduced parenteral glucose supply protected against neonatal sepsis via decreased bacterial burden and dampened pro-inflammatory immune responses. Relative to our previous findings, we showed that a more moderate reduction in glucose intake provided similar positive benefits in outcomes, while avoiding hypoglycemia^27^.

### Reduced glucose supply increases hepatic OXPHOS and gluconeogenesis and attenuates inflammatory pathways

Given that both glucose levels and bacterial burdens were lower in the reduced glucose group we speculated that proliferation of bacteria was driving the phenotype. However, within SE-Low and SE-High pigs, mortality did not correlate to bacterial burdens (Sup. Figure 1F). Although we cannot exclude that circulating glucose levels affected bacterial proliferation, we hypothesized that an important mechanism behind these positive effects was a reduction of hepatic and circulating glycolytic metabolism, which may lead to rewiring immune cells to rely on other substrates than glucose, thereby constraining the overall pro-inflammatory cascades. Thus, we explored how glucose supply affected immune-metabolic gene expression in the liver, collected at euthanasia. Both infection and glucose supply dramatically impacted hepatic gene transcription, with more than 4000 differentially expressed genes (DEGs) for each group comparison (Figure 2A and Sup. Table S1A-D). Relative to SE-High, the SE-Low animals showed elevated pathways related to glycolysis/gluconeogenesis, OXPHOS and TCA cycle, as well as fatty acid and glucogenic amino acid metabolism (Figure 2B, C, Sup. Figure S2 and all DEGs can be found in Sup. Table S1). Though glycolysis and gluconeogenesis share many enzymes catalyzing reversible reactions, enzymes related solely to glycolysis or glycogen formation, such as glucokinase (*GCK*) and phosphoglucomutase 2 (*PGM2*) were downregulated in SE-Low compared to SE-High (Figure 3D, E), whereas those only involved in gluconeogenesis, such as fructose-bisphosphatase 2 (*FBP2*) and pyruvate carboxylase (*PC*) were upregulated (Figure 3F, G). This suggests that a shift from glycolysis to gluconeogenesis occurs in infected animals kept on a reduced glucose supply, even during infections. In parallel, SE-Low animals showed multiple attenuated pathways related to inflammation, including NF-kappa B, IL-17, TNF, and Transforming growth factor-β (TGF-β) signaling (Figure 3C, Sup. Figure S2, Sup. Table S1A). Within the two un-infected control groups, glucose supply affected similar metabolic processes, but not inflammatory pathways (Sup. Table S1B, E-F).

**Figure 2:**
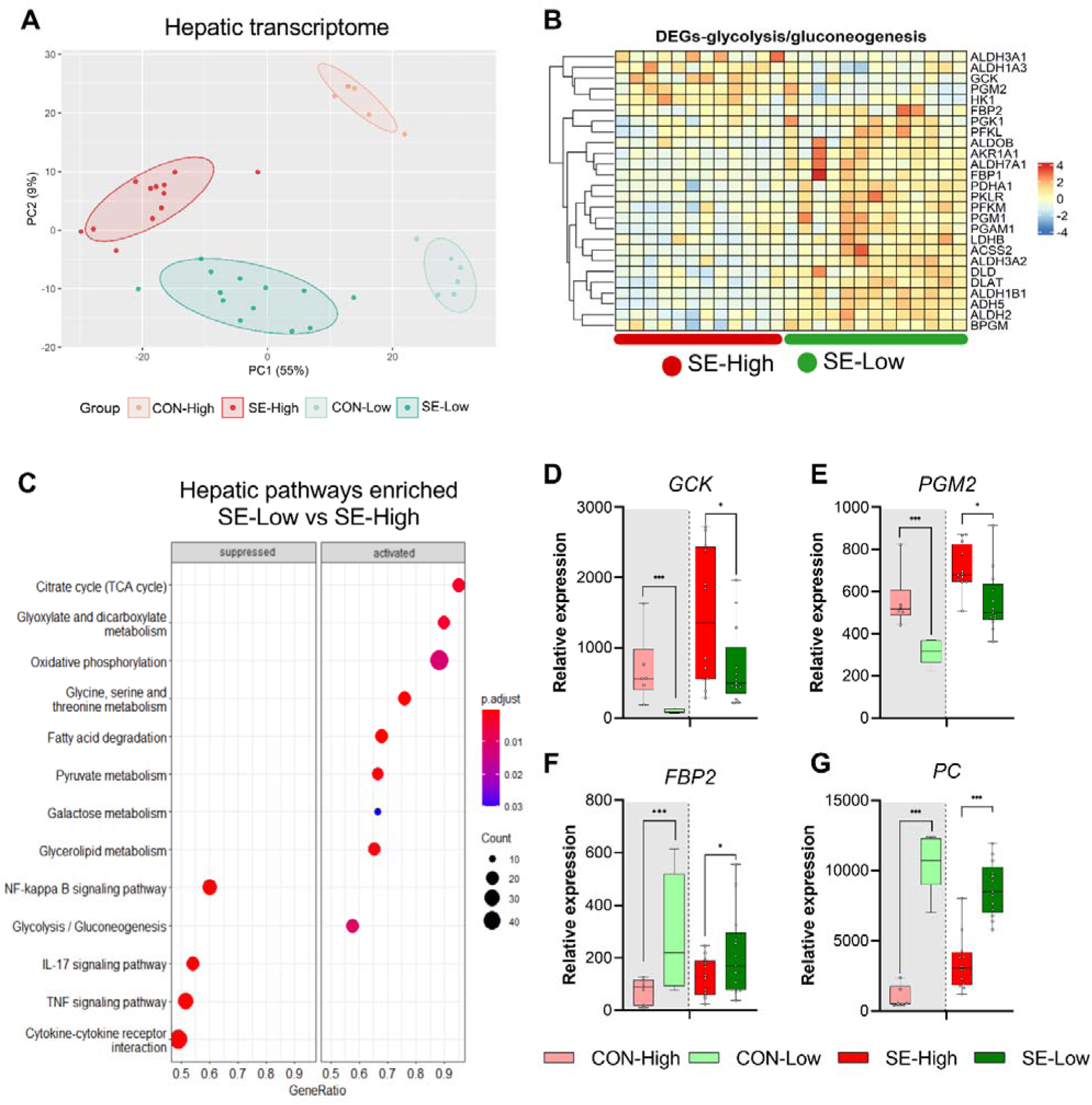
Impacts of glucose supply on liver transcriptomics at euthanasia. **A**: principal component analysis plot of transcriptomic profiles among the four groups. **B:** Heatmap of differentially expressed genes (DEGs) related to glycolysis/gluconeogenesis, in the enriched pathways between the two infected groups. Differences shown as Z-scores, where red color indicates a higher expression and blue a lower. **C**: Gene set enrichment analysis (GSEA) with gene ontology database showing the top pathways activated and suppressed by reduced glucose supply in infected animals. Size of dots indicates number of DEGs while red color indicates lower adjusted P value. **D-G:** Expression of genes exclusively related to glycolysis or gluconeogenesis, shown as relative expressions using 95% box plots. *: P < 0.05, ***: P < 0.001, n = 7-8 for each control group and 15-16 for infected groups.

**Figure 3:**
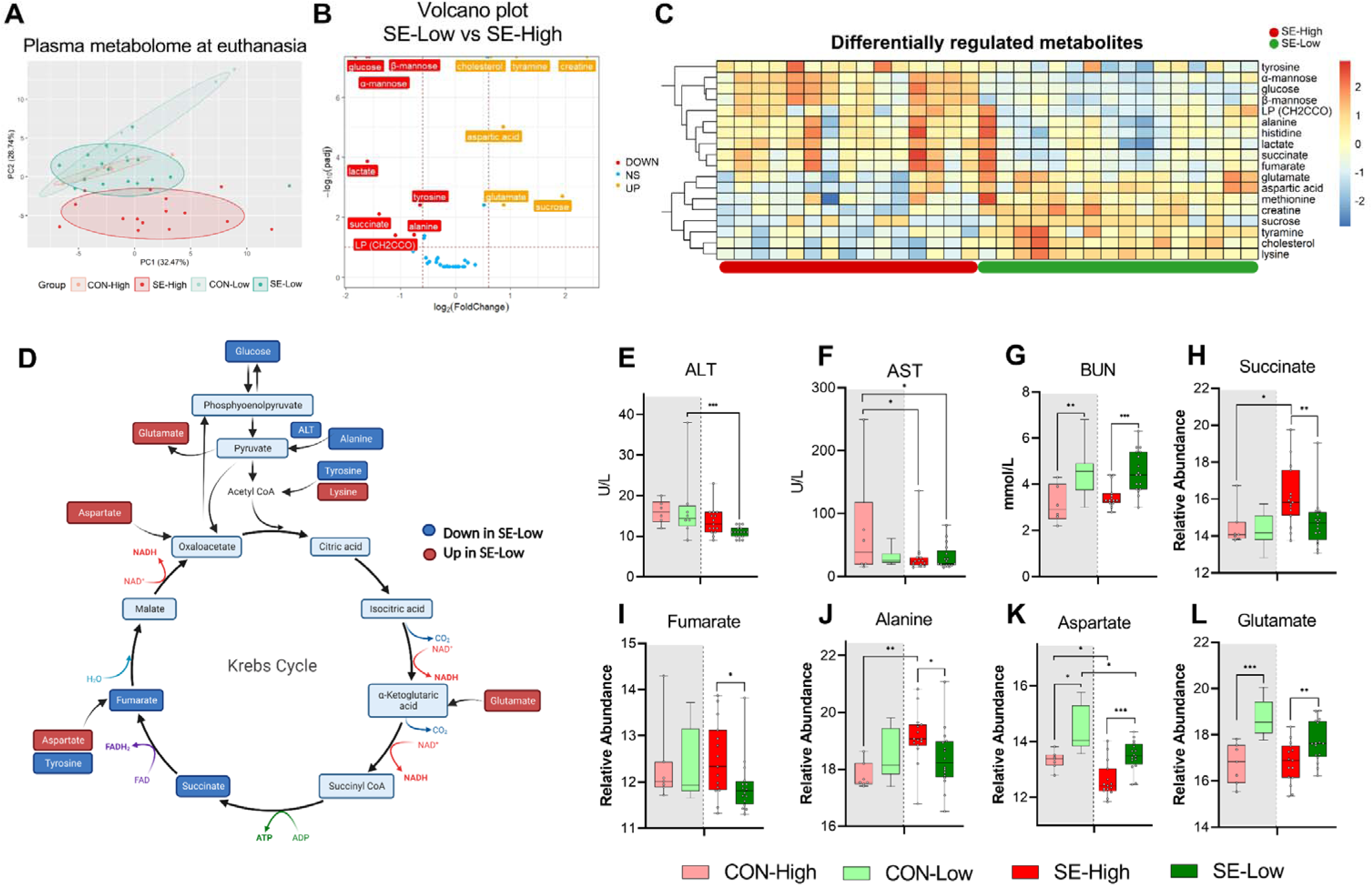
Impacts of glucose supply on plasma metabolism response revealed by proton NMR-based metabolomics at euthanasia, either at humane endpoint or 22 hours after bacterial inoculation. **A:** Score plot of principal component analysis performed on metabolomics data among the four groups. **B:** Volcano plot showing the differential expressed metabolites (DEMs) in SE-Low vs SE-High. Yellow color indicates higher plasma levels, red lower. **C:** Heatmaps of identified DEMs between the two infected groups. Differences shown as Z-scores, where red color indicates a higher expression and blue a lower. **D:** Schematic representation of gluconeogenesis and TCA cycle metabolites altered between the two infected groups. Red represents a metabolite upregulated in SE-Low vs SE-High whereas dark blue indicates a metabolite downregulated; light blue indicates a metabolite not detected by ^1^H NMR. **E-G:** Plasma alanine transaminase (ALT), aspartate transaminase (AST), and BUN levels, shown as 95% box plots. **H-L:** DEMs involved in gluconeogenesis and TCA cycle, shown as 95% box plots. *: P < 0.05, **: P < 0.01, ***: P < 0.001, n = 7-8 for each control group and 15-16 for infected groups.

### Plasma metabolome confirms the glucose impact on gluconeogenesis and balanced TCA cycle

With the profound impact of infection and glucose supply on the transcribed hepatic metabolic pathways, we further sought to analyze the plasma metabolome at the time of euthanasia to gain more insights on these metabolic processes, especially to confirm if enhanced gluconeogenesis occurred in SE-Low animals. Via proton nuclear magnetic resonance (^1^H NMR) spectroscopy, we identified 70 major metabolite signals, among them, a total of 43 plasma metabolites were annotated. The metabolite profile of SE-High animals appeared to be distinct from the remaining three groups (Figure 3A). Infection changed abundance of 11 and 28 metabolites in reduced and high glucose conditions, respectively, while 35 metabolites were differentially regulated between the two infected groups (All differentially expressed metabolites shown in Sup. Table S3). Major annotated differentially regulated metabolites and blood biochemical parameters between SE-Low and SE-High groups were mainly sugars, amino acids and TCA cycle intermediates (Figure 3B,C). Figure 3D highlights the detected metabolomic changes involved in the gluconeogenesis and TCA cycle. Importantly, SE-High animals showed higher levels of succinate and fumarate (Figure 3H-I), further supporting that high glucose supply rewired systemic metabolism to heightened glycolysis, via inhibiting enzymes catalyzing succinate conversion, causing accumulation of succinate and fumarate^32^. On the other hand, SE-Low animals showed decreased plasma alanine and alanine aminotransferase activity, and increased levels of blood urea nitrogen, glutamate and aspartate (Figure 3E, G, J, L), all pointing to the direction of enhanced gluconeogenesis by metabolizing glucogenic amino acids, possibly contributing to the maintenance of glucose homeostasis under reduced glucose supply.

### Only inflammatory but not metabolic network of blood transcriptome is modulated by reduced glucose supply

We then explored how infection and glucose supply affected blood leucocyte gene expressions at 12 h post-infection challenge, the time-point when clinical outcomes started to drastically differ between the two infected groups. Infection itself dramatically altered the whole blood transcriptome (Figure 4A), with more than 5000 differentially expressed genes (DEGs) (all shown in Sup. Table S2). Glucose supply in un-infected conditions had limited impact, while within the infected groups differences in gene expression were much less pronounced than in the liver. The SE-Low animals showed 321 up-regulated and 363 down-regulated DEGs, compared to SE-High animals (Sup. Table S2A). Pathway enrichment analysis for DEGs between the two infected groups showed dampened immune response, enhanced glycerolipid metabolic processes, and re-organization of cellular structure in SE-Low animals (Figure 4B-E, Sup. Figure S3). Surprisingly, no enriched pathways related to the metabolism of glucose, fatty acids or amino acids were observed. These data strongly suggest that modulations of hepatic, but not circulating immune cell metabolism, by reduced glucose supply play a central role in guiding systemic inflammatory responses in preterm neonates.

**Figure 4:**
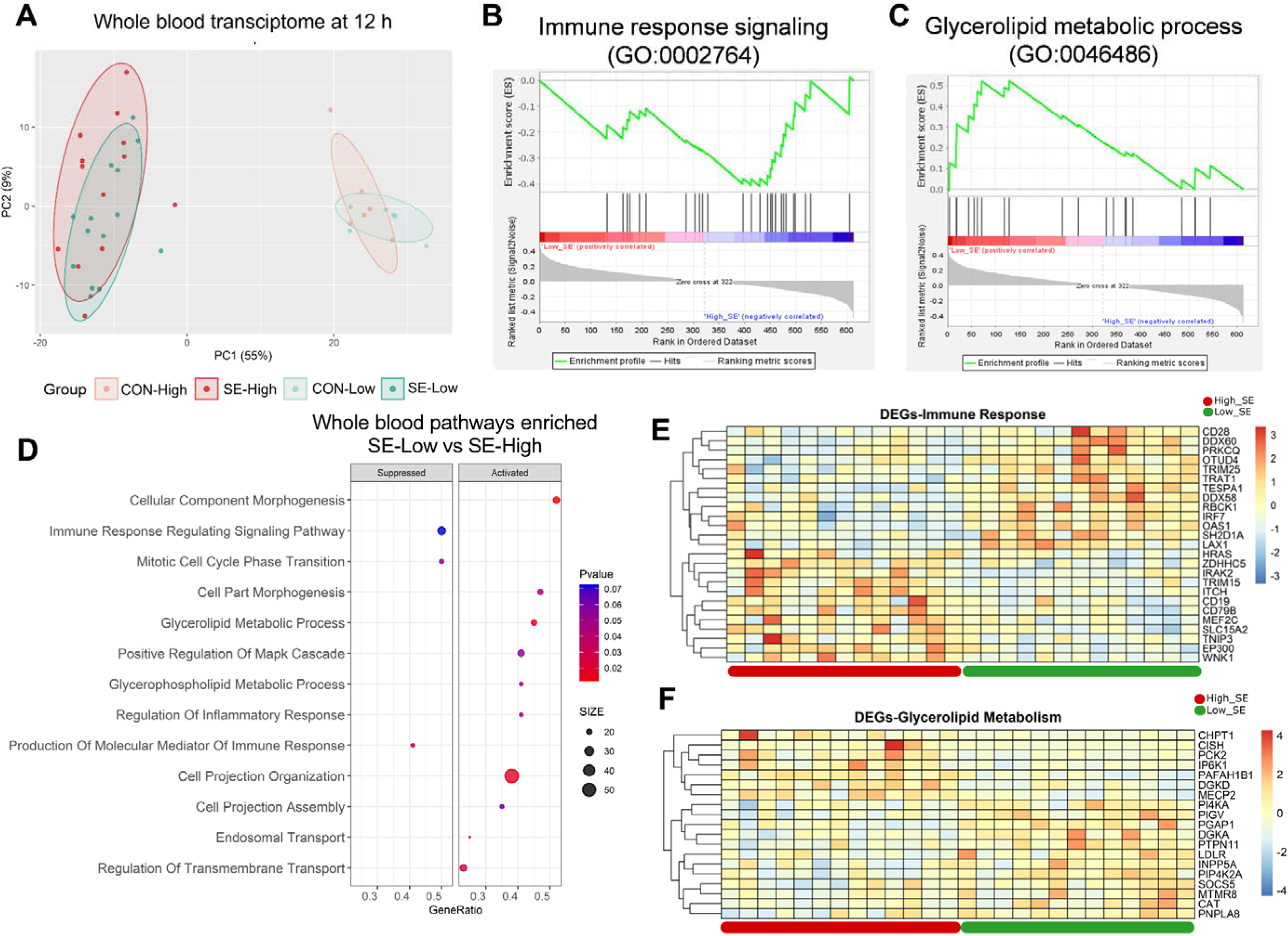
Impacts of glucose supply on plasma proteome at different time points post-bacterial infection. **A-D:** Score plot of principal component analysis of proteome profiling changes from 3 hours to euthanasia among the four groups. **E-F:** Top pathways differing between SE-Low and SE-High at 12 hours post bacterial inoculation or euthanasia, were enriched using DAVID. Involved differential expressed proteins (DEPs) counts displayed on X axis and size of dots indicates number of DEPs, while red color indicates lower P value. **G-I:** DEPs involved in glycolysis or gluconeogenesis, including Lactate Dehydrogenase B (LDHB), Aldolase, Fructose-Bisphosphate A (ALDOA), Dihydrolipoamide dehydrogenase (DLD. **J-L:** DEPs of porcine IAIP related proteins, heavy chains 1 and 2 (ITIH1, ITIH2) and Alpha-1-Microglobulin bikunin precursor (AMBP). **G-L:** Shown as relative abundances with 95% box plots, data analyzed separately for each timepoint, bars labeled with different letters are significantly different from each other (P < 0.05), n = 6 in each group.

### Plasma proteome confirms groups of interacting mediators involved in immune-metabolic modulations

Clinical, transcriptome and metabolome analysis at 12 h post-infection and euthanasia clearly showed distinct immune-metabolic signatures between the two infected groups. To further capture these differences, we performed plasma proteomics on longitudinal samples, from 3 h post-bacterial challenge to euthanasia. Principal component analysis of proteomic profiles (Figure 5A-D) demonstrated no separation among the four groups at 3 h and marginal separation of SE-High from SE-Low at 6 h. The full impact of infection and glucose supply during infection was only obvious at 12 h and euthanasia. These findings are in line with hematological and plasma cytokine data, which revealed only limited changes at 3-6 h and more dramatic differences from 12 h until euthanasia.

**Figure 5:**
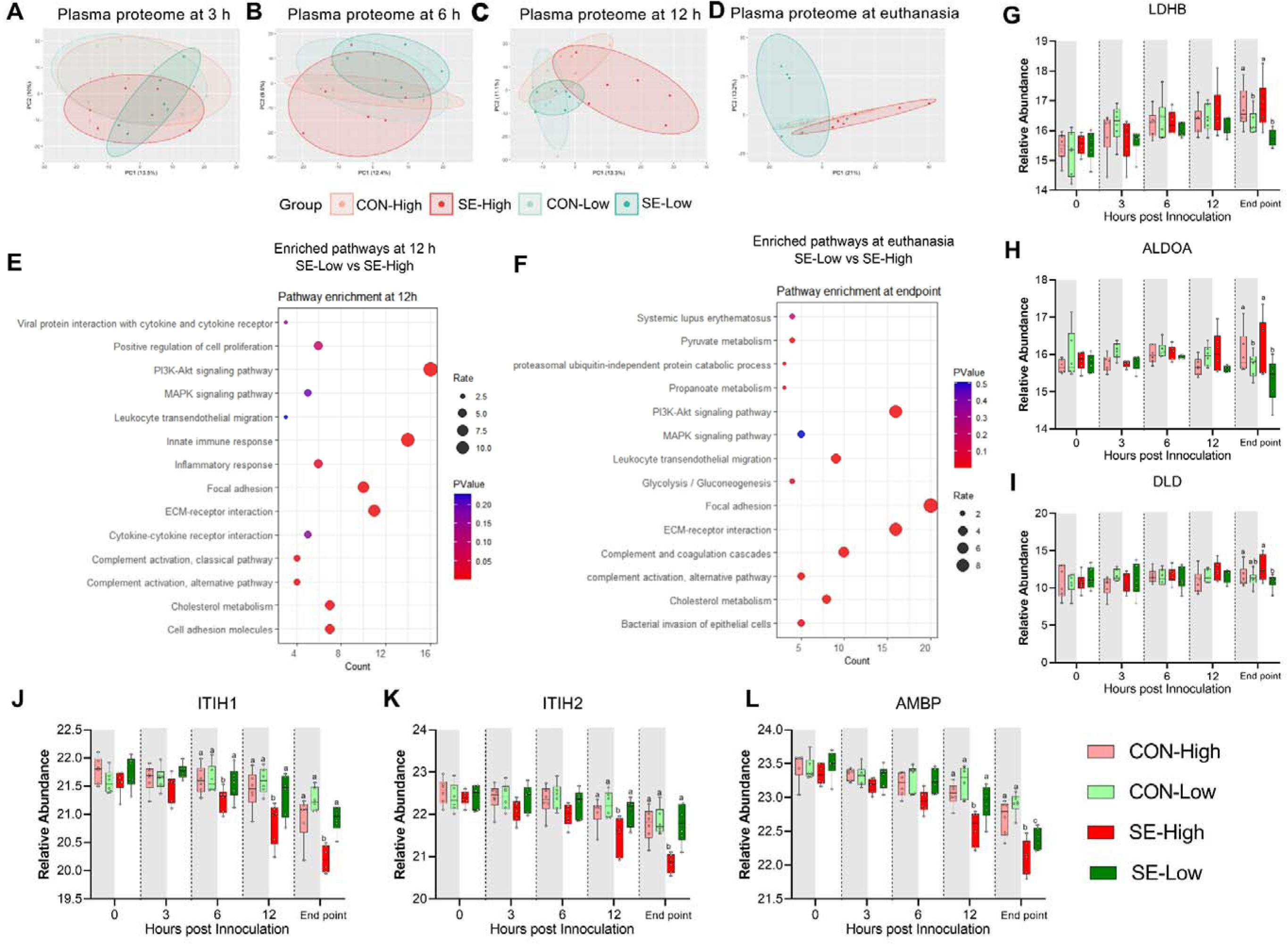
Impacts of glucose supply on whole blood transcriptomics at 12 hours after bacterial inoculation. **A**: Score plot of principal component analysis of transcriptomic profiles performed among the four groups. **B, C:** Gene set enrichment analysis (GSEA) of the immune response signaling (GO:0002764) with negative enrichment score and glycerolipid metabolic process (GO:0046486) with positive enrichment score in the SE-Low, relative to SE-High animals. **D**: Enrichment analyses using GSEA with gene ontology database showing the top pathways activated and suppressed by reduced glucose supply in infected animals. Size of dots indicates number of DEGs while red color indicates lower adjusted P value. **E, F**: Heatmaps of differentially expressed genes (DEGs) involved in the enriched immune response and glycerolipid metabolism pathways between the two infected groups. Differences shown as Z-scores, where red color indicates a higher expression and blue a lower. n= 6 in each control group and 13-14 for infected groups.

Due to relatively low differentially expressed proteins (DEPs) between the two infected groups (93 and 179 proteins at 12 h and euthanasia, respectively), we performed pathway enrichment analysis with all DEPs (Sup. Table S4). We detected similar enriched pathways to other transcriptome/metabolome data at 12 h and euthanasia: immune response, pyruvate metabolism and glycolysis/gluconeogenesis (Figure 5E-F). Specifically, levels of proteins encoding for enzyme conversion from pyruvate to lactate (LDHB, DLD) and upstream conversion of glucose in the glycolysis pathway (ALDOA) were higher in animals receiving high vs. reduced glucose supply, irrespectively of infection status, again showing increased use of glycolysis (Figure 6G-I). To further explore differences in inflammatory pathways we investigated plasma levels of IAIP protein subunits, as these are known to be reduced in infants with bacterial sepsis^16^. Although, at the end of the experiment, AMBP levels were lower in SE-Low compared to the uninfected group (Figure 5L), levels of heavy chains 1 and 2 as well as alpha-1-microglobin precursor to bikunin (AMBP) were significantly higher in SE-Low compared to SE-High, which correlates with a less severe inflammatory response (Figure 5K-L).

**Figure 6:**
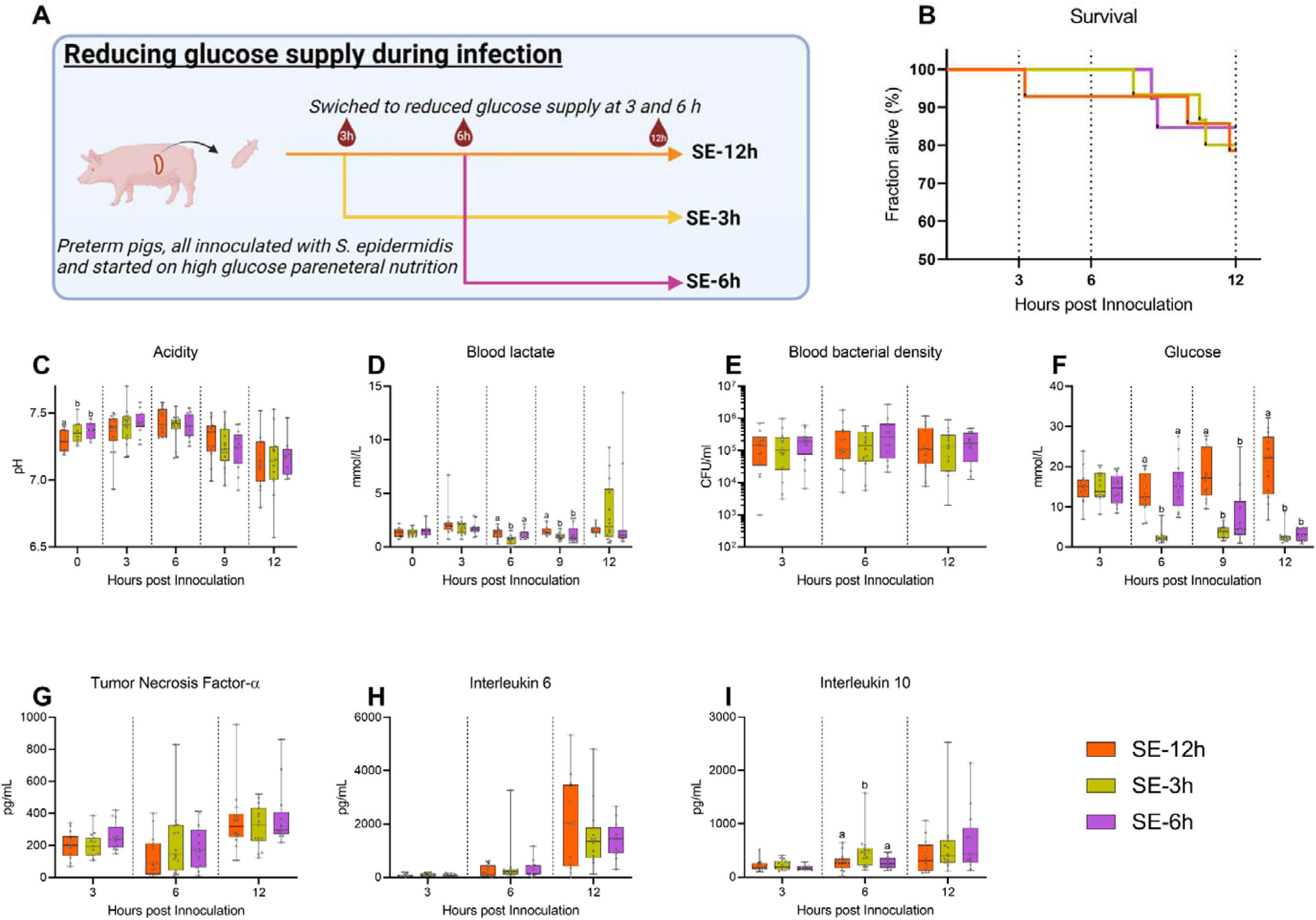
Follow-up experiment investigating effects of changing glucose regimen during infection. **A:** Study overview, preterm pigs were all started on high glucose regimen and inoculated with *S. epidermidis*. After 3 hours one group was shifted to low glucose PN (SE-3h), while after 6 hours another group was similarly shifted (SE-6h) and the remaining pigs continued on high glucose PN for the rest of the experiment (SE-12h). Figure created with BioRender.com. **B:** Survival during experiment, presented as Kaplan-Meier curves. **C, D:** Blood gas data collected at baseline and 3-12 hours after inoculation. **E:** Blood bacterial density 3-12 hours after inoculation, shown on a logarithmic scale. **F:** Blood glucose levels 3-12 hours after inoculation. **G-I:** Plasma cytokine levels 3-12 hours after inoculation. **C-I:** Presented as 95% box plots, data at each time point analyzed separately, bars labeled with different letters are significantly different from each other (P < 0.05), n = 11-14 for SE-12h, n = 12-15 for SE-3h and n = 10-12 for SE-6h.

### Timing of glucose supply guides inflammatory and clinical outcomes post-infection

In a clinical setting, any treatment of infection would not be initiated before clinical symptoms become evident, which in our model occurs 3-6h after bacterial inoculation. We therefore investigated whether an intervention to reduce the high parenteral glucose supply at either 3 or 6 h post inoculation could rescue the animals from sepsis. *S. epidermidis* infected preterm pigs were all started on a high glucose supply and then switched to a reduced glucose supply at either 3 h (SE-3h) or 6 h post-inoculation (SE -6h) or kept on high glucose PN (SE-12h) and followed until 12 h post infection (see Figure 6A for experimental overview).

No difference in survival could be detected among these three infected groups until 12 h (Figure 6B). Likewise, although blood pH gradually dropped, it was similar among the groups 3-12 h post-infection (Figure 6C). However, shifting to a reduced glucose supply led to lower lactate levels at 6 h post infection in SE-3h pigs, while at 9 h the lower levels were observed in both shifted groups (Figure 6D). Although there were no differences in blood bacterial density at any timepoint (Figure 6E), blood glucose levels were quickly affected by shifting to a reduced glucose regime (Figure 6F), while only minor effects on hematological parameters were observed (Sup. Figure S4A-D), with similar plasma TNF-α and IL-6 responses (Figure 6G, H). Plasma IL-10 levels were transiently elevated in the SE-3h group 6 h after inoculation, indicating that some increase in anti-inflammatory responses (Figure 6I). In summary, glucose reduction during infection in this animal model did not improve clinical status until 12 h post challenge. Therefore, it appears that high blood glucose levels at the time of or shortly after the infection, rather than at clinical manifestation, are a determining factor for the clinical fate of infected preterm pigs.

### Administration of hIAIP combined with low glucose PN does not improve septic conditions

Since reduced parenteral glucose regimen during neonatal infection led to improved survival, but not complete protection from sepsis, by modulating the hepatic immune-metabolic networks leading to reduced inflammation: We therefore wished to explore whether concomitant intravenous administration of purified hIAIP, a liver derived protein, could provide additional treatment benefits. Animals were provided with previously reported doses of hIAIP^18^ at one and 12 h and monitored until 22 h post-infection (SE-IAIP; Figure 7A). Supplementation of hIAIP increased mortality from 12 h onwards (Figure 7B), but induced only limited alterations in blood acidity or lactate levels (Figure 7C,D). However, blood bacterial density was higher by the end of the experiment in the SE-IAIP group compared to SE-Low, despite similar glucose levels (Figure 7E,F). Interestingly there was a small, but significant, increase in plasma levels of IL-10 3 h post-infection (Figure 7G). However, plasma levels of IL-6 or TNF-α, as well as blood leucocyte counts did not differ between SE-Low and SE-IAIP animals, although the SE-IAIP group did not recover neutrophils to the same degree as SE-Low by the end of the experiment (Sup. Figure S5A-E). Therefore, in contrast to the hypothesized beneficial complementary effects of hIAIP treatment and low PN glucose regimen, the combined treatment resulted in reduced blood bacterial clearance *in vivo* and decreased survival of the infected animals.

**Figure 7:**
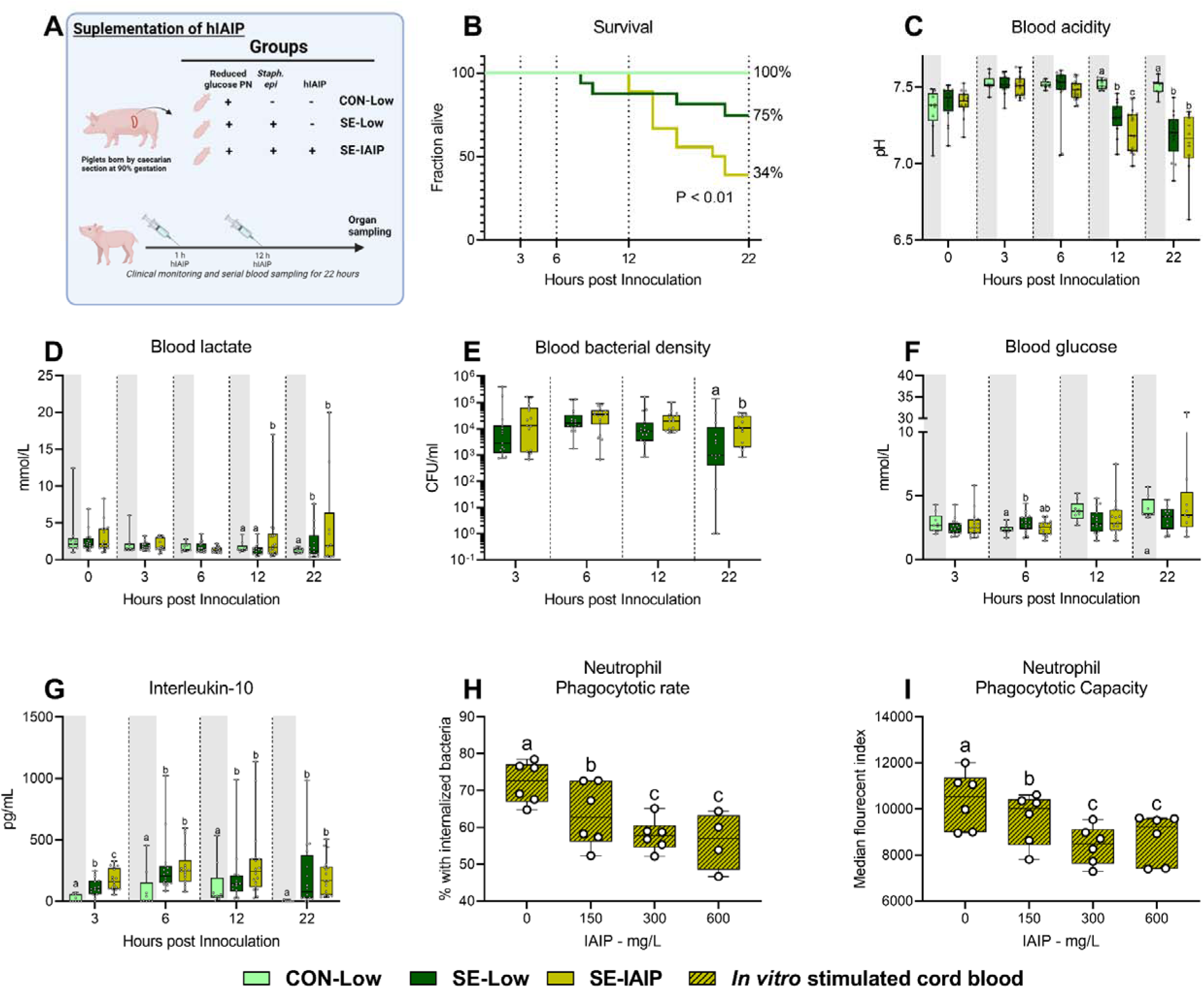
Human IAIP intervention, clinical and immunological results. **A:** Study overview, preterm pigs were nourished low (5%, 7.2 g/kg/d) glucose parenteral nutrition and infused with either *Staphylococcus epidermidis* or control saline. At one and 12 hours post inoculation, infected animals were treated with either human IAIP (50mg/kg) or saline and followed for 22 hours while blood samples collected for further analysis. Figure created with BioRender.com. **B:** Survival of animals during experiment, presented as time to euthanasia according to predefined humane endpoints with corresponding log-rank test comparing SE-Low and SE-IAIP **C:** Blood glucose, measured by glucose meter 3, 6 and 12 hours after SE inoculation as well as at euthanasia, presented as 95% box plots. **D,E:** Blood gas parameters collected before inoculation with SE (0 hours) and 3, 6 and 12 hours after as well as at euthanasia, presented as 95% box plots. **F:** Blood bacterial density in infected groups at 3, 6, 12 and 22 hours after bacterial inoculation, presented as 95% box plots on a logarithmic scale. **G:** Plasma IL-10 and blood neutrophil fraction at 3, 6, 12 and 22 hours after bacterial inoculation, presented as 95% box plots. **H: I:** Cord blood neutrophil phagocytic rate and capacity following *in vitro* challenge with fluorescently labeled E. coli and treatment with increasing doses of IAIP, n = 7. **C-H:** Data at each time point analyzed separately, bars labeled with different letters are significantly different from each other (P < 0.05), n = 8 for control animals, and 10-18 for infected groups.

To further explore the underlying cause of this phenotype, we turned to an *ex vivo* approach, assessing phagocytosis of bacteria by cord blood neutrophils from preterm pigs, treated with increasing hIAIP concentrations. The percentage of phagocytic neutrophils and the number of engulfed bacteria were inversely proportional to the levels of hIAIP received (Figure 7H-I). This demonstrated an inhibitory effect on neutrophil activation *in vitro*, thus suggesting that hIAIP-mediated reduction in neutrophil phagocytic capacity results in increased pathogen load and reversal of effects induced by the reduced glucose regimen.

## Discussion

Using a clinically relevant animal model of infections in preterm neonates, we found that a a reduced parenteral glucose supply led to better survival less clinical deterioration and reduced circulating bacterials, as well as dampened pro-inflammatory responses, all while avoiding hypoglycemia. It was evident that all markers of neonatal sepsis, such as blood pH, lactate, leucocyte/thrombocyte counts were improved by administration of the low glucose PN. At the same time, this nutritional strategy had a clear impact on the hepatic metabolism, reducing the rate of glycolysis while favoring OXPHOS and possibly gluconeogenesis. However, given that most metabolic endpoints are collected at euthanasia we are unsure when the shift in metabolism occurs. Also, to our surprise we found only minor changes in the metabolism of circulating leucocytes despite downregulation of a range of inflammatory genes in the SE-Low group. As such, it is unlikely that the blood leukocyte pool is the crucial link between glucose sensing and immune response to systemic infections. Instead, much of the cytokine production during infections seems to derive from liver macrophages or splenocytes, organs that monitor for blood borne pathogens^33,34^. In the liver transcriptome, there were clear signs of both diminished pro-inflammatory responses and a switch to OXPHOS in SE-Low animals and, when coupling with the plasma metabolome, signs of enhanced gluconeogenesis. We are however limited by the use of whole blood/tissue transcriptomics, as opposed to single cell sequencing, which does not allow for the identification of changes to individual cell types. The observed effects likely occur in separate cell types, with metabolic modulations by glucose supply probably more pronounced in hepatocytes while the immunological changes were altered in the liver immune cells. It is likewise possible that metabolic adaptations in circulating immune cells are present in specific subsets, not detected by the whole blood sequencing. However, from the entirety of our data the liver appears to be the orchestrator of the immune response to bacterial infection in preterm neonates and its immune-metabolic response is affected by the circulating glycemic state.

Crucially, a moderately reduced glucose supply did not lead to severe hypoglycemia as we had observed in the previous study^27^. However, we also found that high glucose provision in the early phases of the infection determined the clinical fate of the animals, as those switched to a reduced glucose regime after the infection was initiated did not have better clinical outcomes than pigs kept on a high-glucose regime throughout the experiment. However, we have not explored the impact of increasing glucose provision after onset of clinical symptoms, so it still unknown if the metabolic state of the immune system at the start of the infection determines the outcome. Interestingly though, given that gluconeogenesis seems to be initiated in the SE-Low group during infection. It is possible that preterm newborns have an adequate capacity to utilize other nutrients (i.e. amino acids) to prevent hypoglycemia during glucose restricted conditions. Therefore, we anticipate that PN with reduced glucose content could be enriched with glucogenic amino acids, which might provide substrates for hepatic gluconeogenesis and help better balance blood sugar levels during neonatal infection.

Reducing glucose intake did not completely protect against development of sepsis, but combining the low glucose regime with supplementation of hIAIP did further not resolve the inflammatory response and failed to improve survival. A possible underlying cause might be the dose used and its known impact on immune cell activation, which has been reported previously^35^. We speculate that in our model of neonatal sepsis, exogenously supplemented hIAIP may have negatively altered responsiveness of neutrophils, thereby potentially increasing pathogen load, resulting in worsening of the septic condition. Likewise, the premature state of the animals, with already suppressed pro-inflammatory responses^36,37^, may have negated the positive results reported elsewhere. Interestingly we observed that porcine IAIP components ITIH1 and bikunin were significantly reduced in circulation only under high PN glucose regimen. This suggests that a high glucose feeding regimen might be a more suitable, alternative study setup to explore potential benefits of IAIP supplementation. Further experiments will be needed to optimize the timing and the context of IAIP administration within the current treatment paradigm to uncover its potential treatment benefits in neonatal sepsis.

In conclusion, we identified a clinically relevant regime of reduced glucose provision to prevent sepsis in infected newborn preterm animals, together with relevant mechanistic insights in hepatic immunity and glucose metabolism. Our findings imply that a similar approach for infants with a high risk of infection may help prevent or attenuate the development of life-threatening sepsis. Currently, we are developing a pilot randomized controlled trial to test the feasibility of such a reduced glucose supply in preterm infants with suspected infection, in parallel with more mechanistic pre-clinical studies of modulated hepatic metabolism and systemic immunity in infected preterm animals. Metabolic modulation of immune responses appears to be a promising therapeutic approach for infections in weak and immunocompromised infants.

## Materials and Methods

### *S. epidermidis* culture preparation

The full procedure for preparation of the *S. epidermidis* bacteria has been described in detail elsewhere^26^. In brief, *S. epidermidis* bacteria from frozen stock were cultured in tryptic soy broth overnight whereafter the bacterial density was determined by spectroscopy and a working solution obtained by diluting with appropriate amounts of sterile saline water.

### Human IAIP preparation

Human IAIP was extracted from fresh frozen human plasma. Starting sample was an intermediate from typical human plasma fractionation process to purify therapeutically relevant plasma proteins, which was first subjected to solvent/detergent treatment using 1% (v/v) Polysorbate 80 and 0.3% (v/v) Tri-n-butyl phosphate. Next, sample was purified by three chromatography steps, comprising anion-exchange chromatography on TOYOPEARL GigaCap Q-650M (Tosoh Bioscience) column, followed by two polishing steps using affinity chromatography on a proprietary synthetic chemical ligand support and hydrophobic chromatography on Phenyl PuraBead®HF (both Astrea Bioseparations). Obtained hIAIPs were further concentrated, yielding a preparation with purity >80%.

#### In vivo procedures, interventions and euthanasia

Preterm pigs (Yorkshire x Duroc x Landrace) were delivered by caesarian section at 90% gestation and rapidly transferred to individual heated, oxygenated incubators. Piglets with prolonged apnea were treated with continuous positive airway pressure until spontaneous respiration was established. Once stable, and still under the influence of maternal anesthesia, all pigs were fitted with an umbilical arterial catheter allowing for the administration of PN and collection of arterial blood samples. Approximately two hours after birth, pigs were stratified by birthweight and sex and, in *Exp 1A*, randomly allocated to receive PN with either high (High, 21%, 30 g/kg/day) or reduced (Low, 5%, 7.2 g/kg/day) glucose levels. After initiation of PN, pigs in each group were further randomly allocated to receive either live *S. epidermidis* or control saline (CON) resulting in four groups: SE-High (n=16), SE-Low (n=16), CON-High (n=8) and CON-Low (n=9). For experimental overview see Figure 1A. Alongside the glucose intervention a separate group of preterm pigs were kept on reduced glucose PN, inoculated with *Staph. Epidermidis* and infused with hIAIP (50 mg/kg, SE-IAIP, n=18) one and 12 h after start of the experiment. These animals were compared to SE-Low and CON-Low pigs described above separately from the effect of the reduced glucose intervention (see Figure 7A).

In a separate follow-up experiment, 41 preterm pigs delivered in the same manner as described above, all started in high glucose PN, were inoculated with *S. epidermidis* and followed for 12h. Pigs were randomly allocated into three groups; with animals either receiving the same high PN glucose supply throughout the whole experiment (SE-12h, n=14),or switched to low glucose supply at precisely 3 h (SE-3h, n=15) or 6 h (SE-6h, n=12) post-infection (See Figure 6A).

Both experiments were initiated by the administration of *S. epidermidis* (1*10^9^ CFU/kg, 0 h) or corresponding volume saline, given as an interatrial infusion over 3 minutes, whereafter pigs were continuously monitored until 12 or 22 h post-infection. During experiments, blood samples were collected through the arterial catheter for blood gas, hematology, and cytokine measurements. For blood bacterial enumeration and glucose measurements, blood was collected by jugular vein puncture at the same timepoints. Blood glucose was measured by capillary puncture using a glucose meter (Accu-Chek, Roche diagnostics, Denmark). At euthanasia, either scheduled or due to clinical deterioration, animals were sedated and euthanized by intracardial injection of pentobarbital after blood had been collected by intracardial puncture to yield plasma and serum. Liver samples were then collected and quickly frozen in liquid nitrogen until further analysis.

#### Humane endpoints for euthanasia

Across all experiments animals were continuously monitored for signs of sepsis. These included: changes in respiratory rate, pallor, increased capillary response time, petechial bleeding, pain, bradycardia and deep lethargy. A blood gas analysis was done on animals showing signs of respiratory or circulatory failure and a pH of less than 7.1 was considered a criterion for immediate euthanasia.

#### Inflammatory markers and biochemistry

Plasma samples collected at various time points after bacterial inoculation were analyzed for porcine specific cytokines using enzyme-linked immunosorbent assay (ELISA), TNF-α (DY690B), IL-10 (DY693B) and IL-6 (DY686, all porcine DuoSet, R&D systems, Abingdon, UK). Plasma samples at planned euthanasia or human endpoints were used for biochemical analysis, using by an Adiva 2120 system (Siemens Healthcare Diagnostics, USA).

#### Blood and liver transcriptomics

Total RNA from whole blood collected at 12 h post-bacterial inoculation and liver at euthanasia was extracted using either MagMAX 96 Blood or Total RNA Isolation Kits (Thermofisher, Waltham, MA) for whole transcriptome shotgun sequencing. Briefly, RNA-seq libraries were constructed using 1000 ng RNA and VAHTS mRNA-seq V3 Library Prep Kit for Illumina (Vazyme, China), and 150 bp paired-end reads of the library were generated using the Illumina Hiseq X Ten platform (Illumina, USA). Quality and adapter trimming of raw reads were performed using TrimGalore (Babraham Bioinformatics, UK), and the remaining clean reads (∼26 million per sample) were aligned to the porcine genome (Sscrofa11.1) using HISAT2^38^. The annotated gene information of porcine genome was obtained from Ensembl (release 99). The script htseq-count^39^ was adopted to generate gene count matrix, followed by analyses of differentially expressed genes (DEGs) using DESeq2^40^.

#### Plasma metabolome

Plasma samples at euthanasia or humane endpoints were subjected to proton (^1^H) NMR spectroscopy-based analysis, as previously described^41^. Briefly, samples were mixed with an equal volume of phosphate buffer solution (pH 7.4, 50 mM) prior to the analysis. ^1^H NMR spectra were recorded on a Bruker Avance III 600 MHz NMR spectrometer (Bruker Biospin, Rheinstetten, Germany) equipped with a 5 mm broadband inverse RT (BBI) probe, automated tuning and matching accessory (ATMA), and a cooling unit BCU-05 and an automated sample changer (SampleJet, Bruker Biospin, Rheinstetten, Germany). The spectra were acquired using a standard pulse sequence with water suppression (Bruker pulse program library noesygppr1d), automatically phase- and baseline-corrected using TOPSPIN 3.5 PL6 (Bruker BioSpin, Rheinstetten, Germany), and referenced to the 3-(trimethylsilyl)propionic-2,2,3,3-D4 acid sodium salt (TSP) signal at 0.0 ppm. The final spectra were analyzed using the Signature Mapping software (SigMa) for identification and quantification of metabolites^42^.

#### Plasma proteome

A subset of plasma samples with sufficient amount from all 5 timepoints of *Experiment 1* (n=6 from each group, from baseline, 3, 6, 12, and 22 hours post-inoculation) were randomly selected for proteome profiling with shotgun LS-MS/MS based proteomics. Briefly, plasma was lysed in cold lysis buffer (5% sodium deoxycholate, 50 mM triethyl ammonium bicarbonate, pH 8.5, Sigma-Aldrich) and prepared with filter-aided sample preparation (FASP). Thereafter proteins were recovered using 10 kDa spin filters (VWR), and samples were reduced, alkylated, and digested with trypsin at 37 °C overnight. Peptides were desalted, dried and stored at −80 °C. At the time of LC-MS/MS analysis, dried peptides were reconstituted in loading buffer (1% acetonitrile and 0.1% formic acid) and determined peptide concentration. Peptides (1µg) were loaded onto a reversed phase column on a Thermo Scientific™ EASY-nLC™ 1200 nanoliquid chromatography system, connected to a Thermo Scientific™ Q Exactive™ HF-X mass spectrometer equipped with a Nanospray Flex™ ion source. A hybrid spectral library was created from DDA (data dependent analysis) and DIA (data independent analysis) search results. For DDA, a modified TOP15 method was adopted to cover the full range MS1 scan with m/z range of 330-1650, followed by 15 data dependent MS2 scans^43^. Data were searched using SpectroMine software (Biognosys, version 2.8), using the Uniprot *Sus scrofa* library from UniProt with the set false discovery rate (FDR) of 1 % and maximal missed cleavages of 2. For DIA, the method consisted of one full range MS1 scan and 21 DIA segments to search peptides using Spectronaut software with FDR set to 1%, as previously described^44^.

#### Effect of human IAIP on in vitro neutrophil function

In addition to the animal study, an *in vitro* experiment to investigate the effect of hIAIP on phagocytic function of neutrophils was performed. Whole cord blood samples from preterm pigs, delivered in the same manner as described above (n=8), were divided and treated with increasing levels of hIAIP (0-600 mg/L, mimicking normal plasma levels in termborn infants^16^) and a standard dose of fluorescently marked *Escherichia coli* (pH Rhodo, Thermofischer) for 30 minutes at 37°C. Samples were then washed, run on a flow cytometer and the neutrophil population was identified, as described in detail elsewhere^45^. Neutrophil phagocytic rate was defined as the fraction of neutrophils with internalized bacteria and the phagocytic capacity as their median fluorescent intensity.

#### Statistics

Data on survival and clinical outcomes were analyzed using Stata 14 (StataCorp, USA). Survival curves were compared using the log-rank test while data collected during the studies were analyzed separately at each timepoint with a two-way ANOVA, with litter and sex as covariates, hereafter group differences were identified with post hoc testing using Tukey’s test. If necessary, data were logarithmically transformed to obtain normal distribution; data that could not conform to normality were analyzed by a non-parametric test.

Omics data were analysed using R studio 4.1.2 (R Studio, Boston, MA). Metabolome and proteome data were analysed by a linear mixed-effect model with treatment as a fixed factor and litter as a random factor at each time point, followed by Tukey Post-hoc pair-wise comparisons using *lme4* and *multcomp* packages^46^. For transcriptomics, significant DEGs among groups were identified by DESeq2 using Benjamin-Hochberg (BH)-adjusted P-value <0.1 as the cut-off. To control type I error, p values were further adjusted by FDR (α = 0.1) into q values^47^. An FDR adjusted P-value < 0.1 for blood and FDR adjusted P-value <0.05 for liver were regarded as statistically significant. Representative plots of single protein or metabolite were presented as violin dot plots with median and interquartile range. All reported measures were evaluated for normal distribution, and logarithmic transformation was performed if necessary.

Pathway enrichment analysis for metabolome data is generated with MetaboAnalyst 5.0^48^ and illustrated with BioRender. For proteomics, gene ontology (GO) and KEGG pathway enrichment analyses for DEPs were performed using DAVID^49^ and a BH-adjusted P-value <0.05 was considered statistically significant. For transcriptomics, gene set enrichment analysis (GSEA) for DEGs was performed using the *fgsea* package with R and GSEA 4.2.2 (UC San Diego and Broad Institute) using GO, KEGG and Hallmark database, and pathways with adjusted P-value <0.05 were considered statistically significant^50^. Oimcs datasets were scaling to unit variance prior to PCA. PCA score plots were generated and the number of principal components was determined by cross validation using R *pcaMethods and prcomp* packages and plotted with ggplot2 package, and heatmaps were generated using R package *pheatmap*.

### Study approval

The animal studies and experimental procedures were approved by the Danish Animal Experiments Inspectorate (license no. 2020-15-0201-00520), which complies with the EU Directive 2010/63 (legislation for the use of animals in research).

## Acknowledgements

The authors thank Thomas Thymann, Simone Offersen, Karoline Aasmul-Olsen, Malene Spiegelhauer, Kristina Larsen, Britta Karlsson and Jane C. Povlsen for the assistance in animal experiments and omic data acquirement and Lucia Gnauer for provision of the purified human IAIP material.

## Author contributions

DNN, IB and PTS designed the experiments while OB contributed to the design. OB, ZW, AB, and DNN performed animal experiments. OB, TM, YX, BK performed laboratory analyses, AMC contributed to proteomic analysis. TM was responsible for bioinformatics under DNN’s supervision. OB and TM performed statistical analysis, managed raw data and generated all figures and tables. OB, IB, PTS and DNN were mainly responsible for data interpretation. OB, TM, and DNN wrote the first manuscript draft. All authors contributed to data interpretation, manuscript revision and approval of the final manuscript version. The levels of contribution in study design, actual experiments, laboratory analyses and writing were used to assign co-first authorship order.

## CONFLICT OF INTEREST STATEMENTS

The study was supported by internal funding at University of Copenhagen and Novo Nordisk Foundation (grant held by DNN, #NNF22OC0078747). The hIAIP intervention was designed, planned, interpreted and financially supported in collaboration with Takeda Pharmaceuticals, division of Plasma-Derived Therapies, Vienna, Austria. Takeda pharmaceuticals assisted in the interpretation of results related to IAIP and did not hold final editorial rights. DNN and PTS were principal investigators on the trial and thus received the funding from Takeda Pharmaceuticals. AMC is an employee of Baxalta Innovations GmbH, a Takeda company. IB is an employee and Baxalta Innovations GmbH, a Takeda company and owns Takeda shares. The other authors declare no conflicts of interest.

## Notes

### Summary of Updates

Revision done following review at eLife Result language updated for clarity. New Sup Fig 1F added shown bacterial burdens over time in animals developing sepsis or not. Discussion language updated for clarity and addition of additional limitations

