## Supplementary material for "Altered hepatic metabolism mediates sepsis preventive effects of reduced glucose supply in infected preterm newborns": Sup. Figures

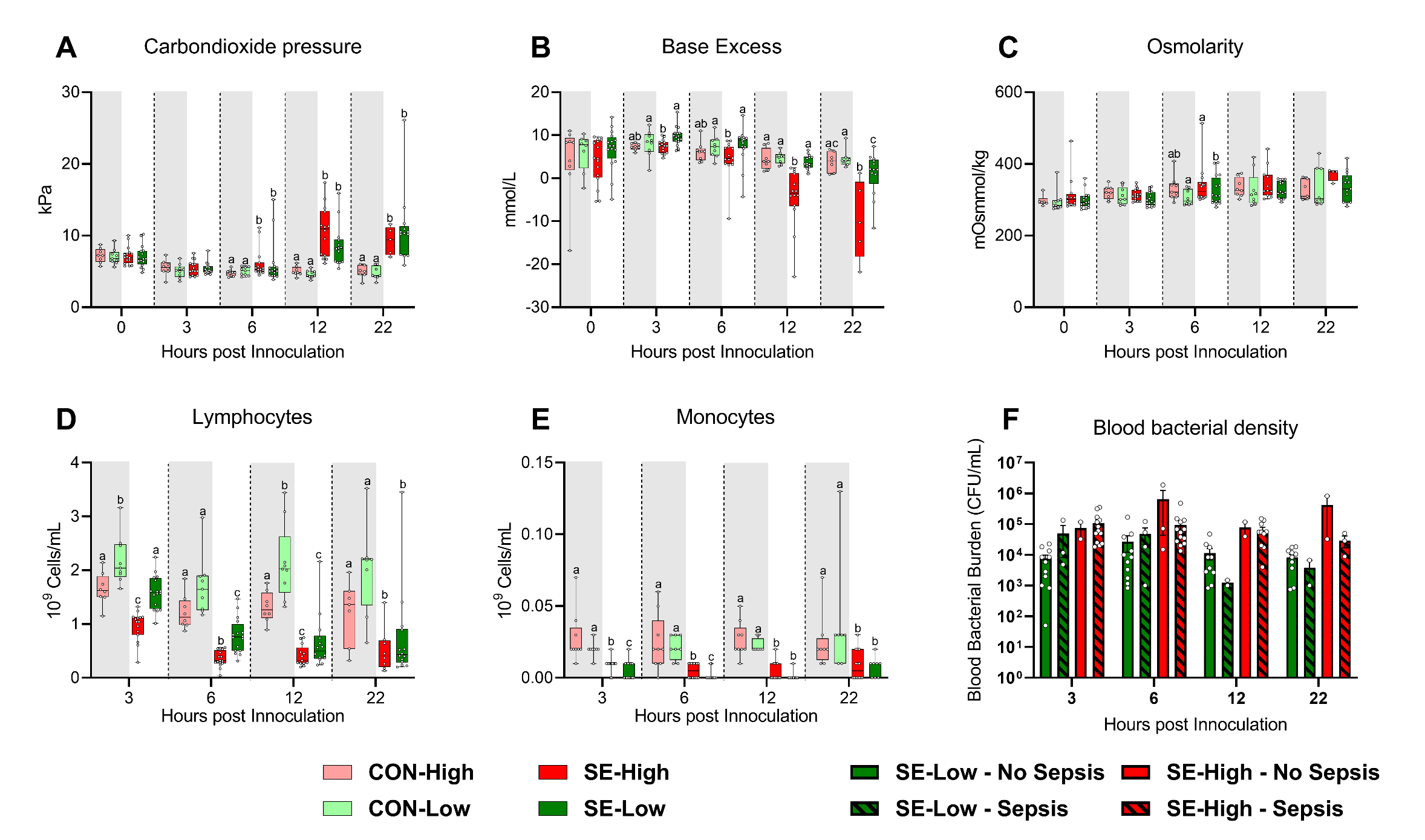

**Sup Figure S1:** Addition clinical results. **A-C:** Blood gas parameters at baseline. **D-E:** Blood hematology 3, 6, 12 and 22 hours after bacterial inoculation. **F:** Blood bacterial burdens in animals that develop sepsis (Lined bars) or not (solid bars) following bacterial inoculation. **A-F:** Data at each time point analyzed separately, presented as 95% box plots or bar charts with standard error, with denoted different letters indicating statistical significances (P < 0.05), n = 8-9 for control animals, and 10-16 for infected groups.

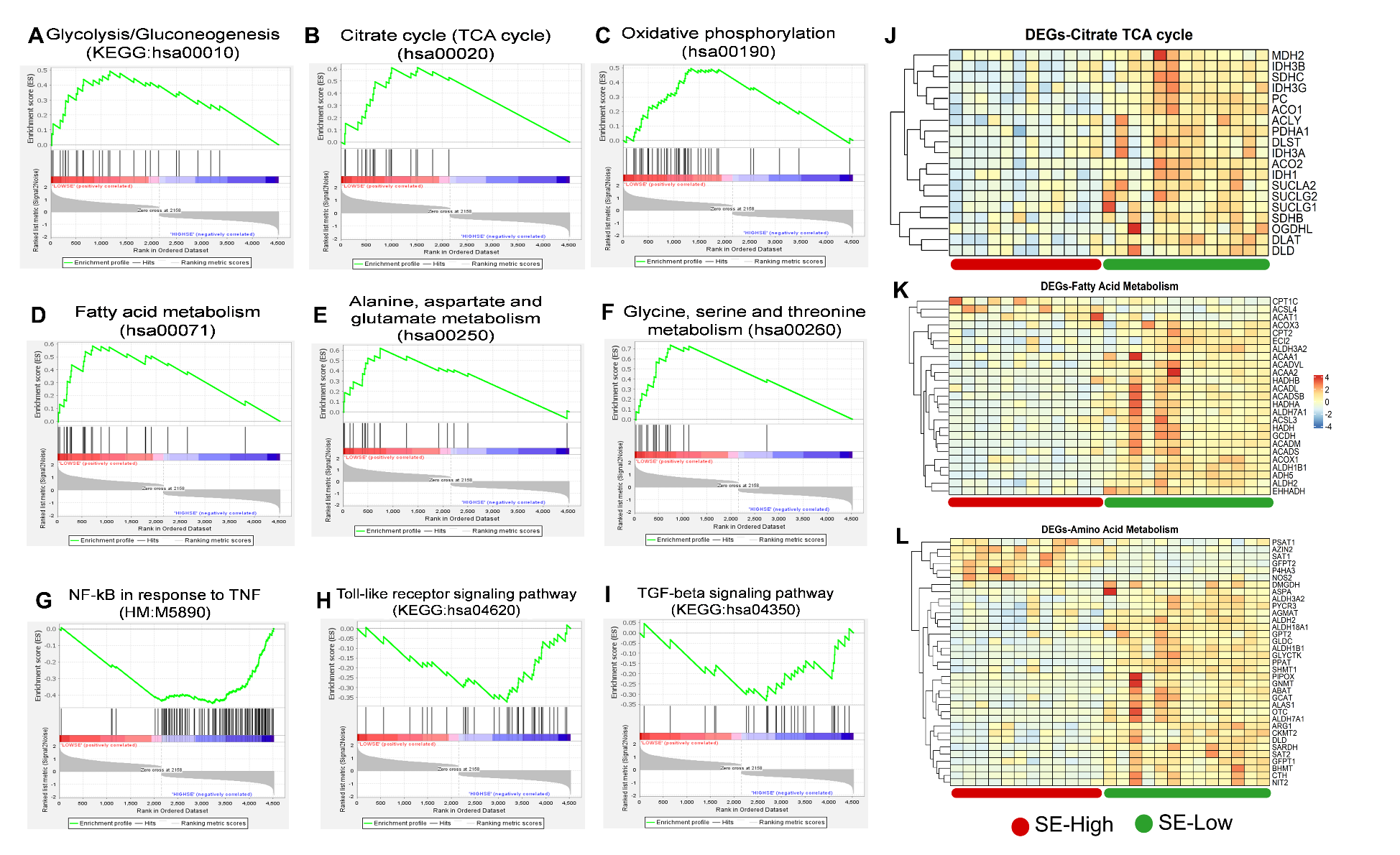

**Sup. Figure S2:** Impacts of glucose supply on inflammation and immune response revealed by liver transcriptomics at euthanasia, either at humane endpoint or 22 hours after bacterial inoculation **A-F:** Gene set enrichment analysis (GSEA) of the energy metabolism related pathways including Glycolysis/Gluconeogenesis (hsa00010), Citrate cycle (TCA cycle) (hsa00020), Oxidative phosphorylation (hsa00190), Fatty acid metabolism (hsa00071), Alanine, aspartate and glutamate metabolism (hsa00250), and Glycine, serine and threonine metabolism (hsa00260) with positive enrichment score in the SE-Low, relative to SE-High animals. **G-I**: GSEA of the inflammation and immune response related pathways using Hallmark and KEGG pathway database, including Genes regulated by NF-kB in response to TNF (M5890), Toll-like receptor signaling pathway (hsa04620), and TGF-beta signaling pathway (hsa04350) with negative enrichment score in the SE-Low, relative to SE-Low animals. **J**-**L:** Heatmaps of differentially expressed genes (DEGs) involved in TCA cycle, fatty acid and amino acid metabolism, in the enriched pathways. Differences shown as Z-scores, where red color indicates a higher expression in SE-Low vs SE-High and blue a lower. N = 15-16 for each group

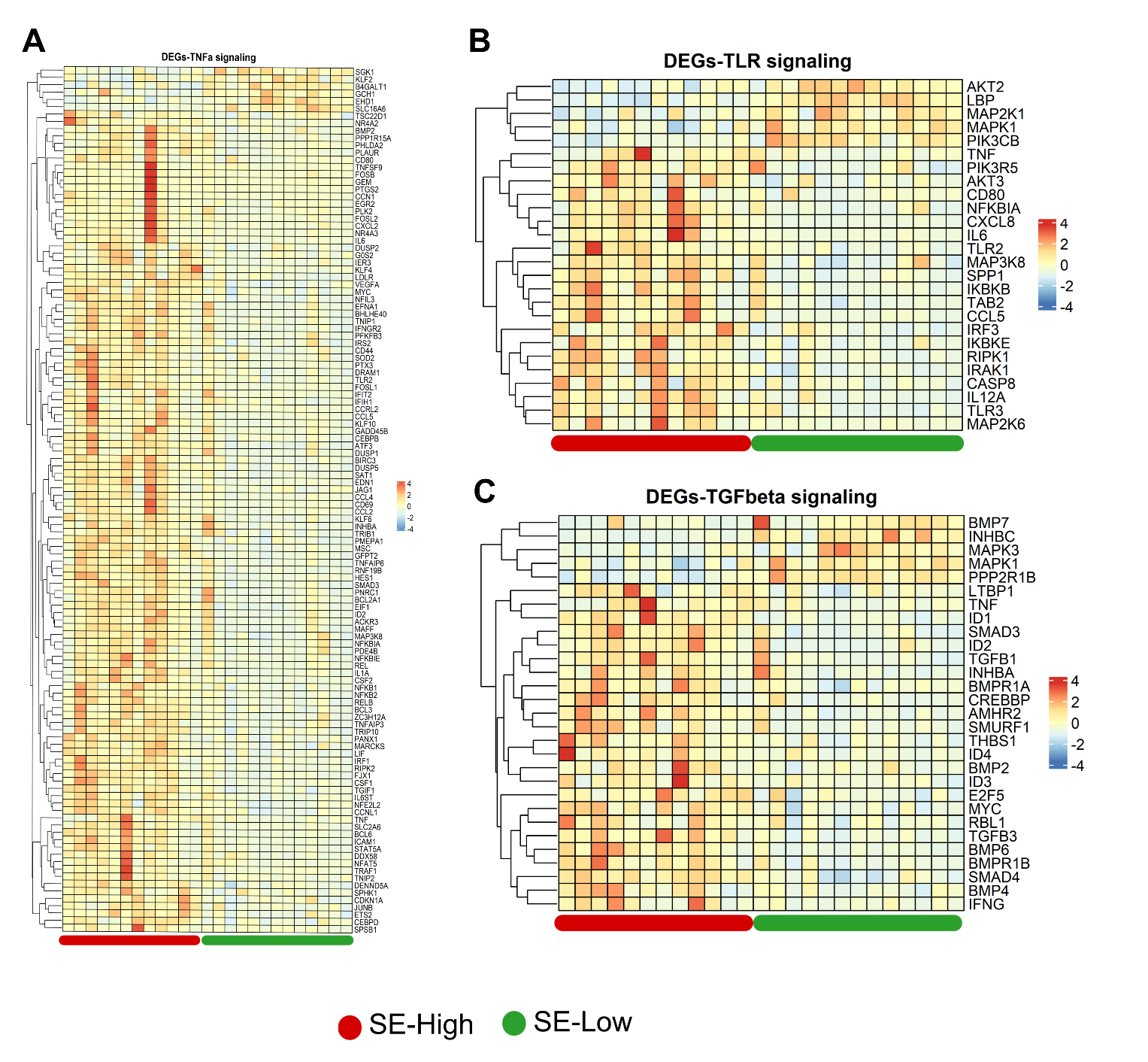

**Sup. Figure S3:** Impacts of glucose supply on whole blood transcriptomics at 12 hours after bacterial inoculation. **A**-**C:** Heatmaps of differentially expressed genes (DEGs) involved in tumor necrosis factor alpha (TNF-α), toll like receptor (TLR) and transforming growth factor beta (TGF-β) signaling in the enriched pathways. Differences shown as Z-scores, where red color indicates a higher expression in SE-Low vs SE-High and blue a lower. N = 15-16 for each group

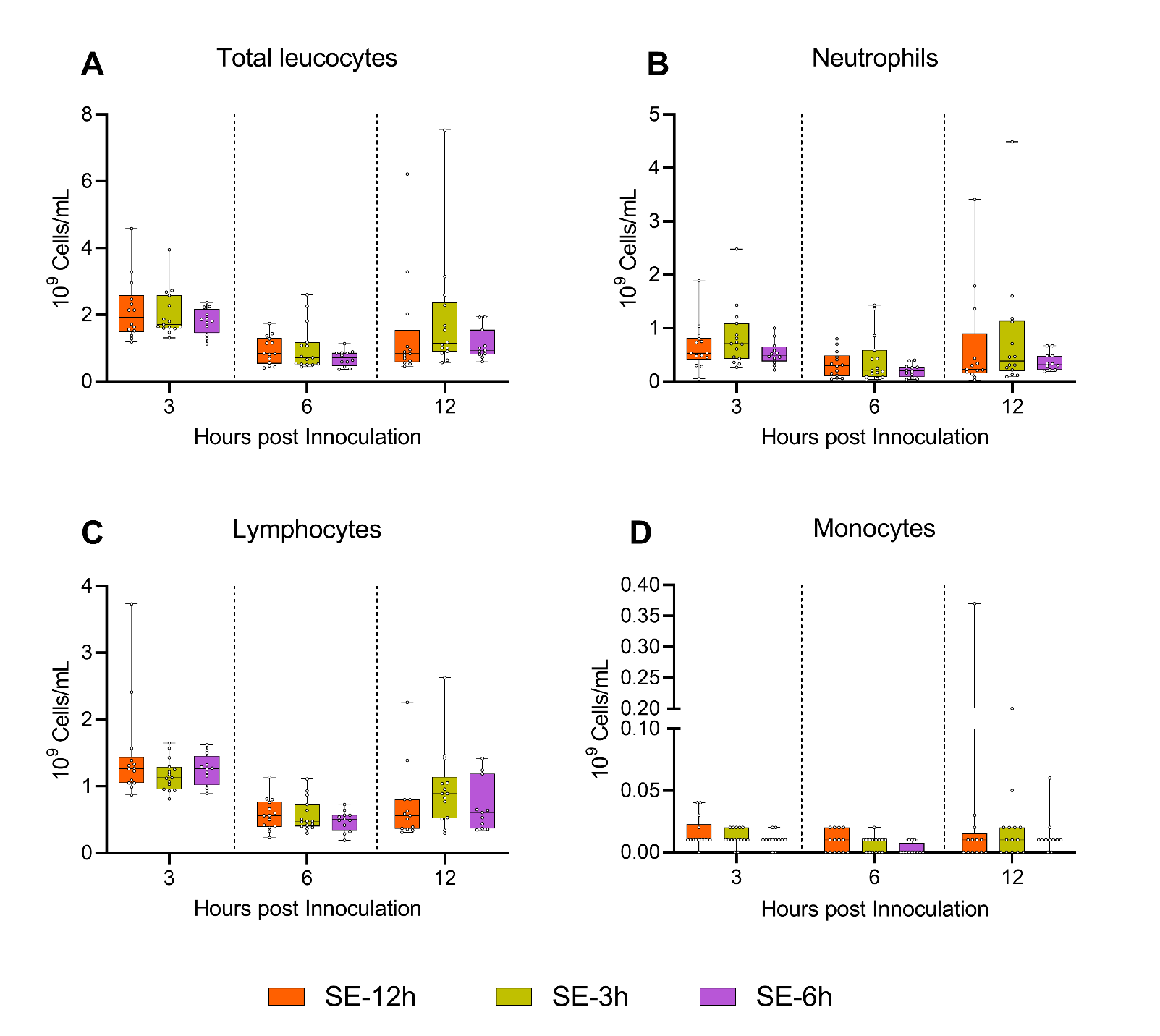

**Sup. Figure S4:** Blood leukocyte subsets during follow-up experiment**. A-D:** Leukocyte subsets at 3, 6 and 12 h after bacterial inoculation, shown as 95% box plots. N = 11-14 for SE-12h, n = 12-15 for SE-3h and n = 10-12 for SE-6h.

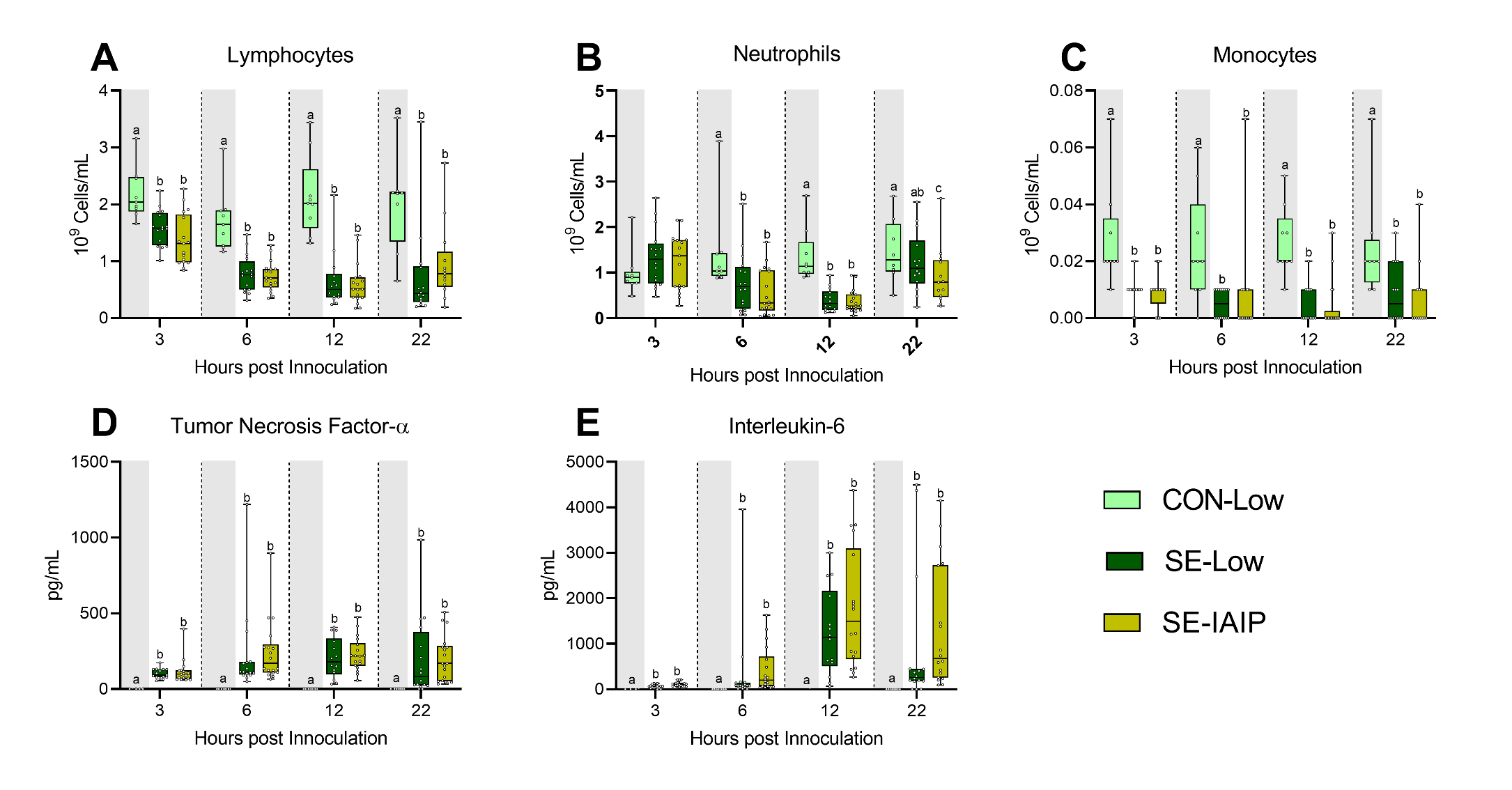

**Sup. Figure S5:** Additional data from human IAIP intervention. **A-C:** Blood leukocyte subsets at 3, 6 and 12 h after bacterial inoculation. D, E: Plasma levels of cytokines at 3, 6 and 12 h after bacterial inoculation. All shown as Data at each time point shown as 95% box plots and analyzed separately, bars labeled with different letters are significantly different from each other (P < 0.05), n = 8 for control animals, and 10-18 for infected groups.
